# End-to-end multimodal deep learning for real-time decoding of months-long neural activity from the same cells

**DOI:** 10.1101/2024.10.14.618046

**Authors:** Yichun He, Arnau Marin-Llobet, Hao Sheng, Ren Liu, Jia Liu

## Abstract

Long-term, stable, and real-time decoding of behavior-dependent neural dynamics from the same cells is critical for brain-computer interfaces (BCIs) and for understanding neural evolution during learning, and memory, and disease progression. Recent advances in flexible and high-density electrodes have enabled the stability required for long-term tracking but generate vast datasets that challenge existing analysis methods. Current spike sorting approaches rely heavily on manual curation and lack scalability for large-scale, real-time processing. Here, we introduce AutoSort, an end-to-end multimodal deep neural network-based method that enables real-time tracking and decoding of the same neurons over months. AutoSort uses a scalable strategy by learning deep representations from initial recordings and applying the trained model in real-time. It integrates multimodal features, including waveform features, distribution patterns, and inferred neuron spatial locations, to ensure robustness and accuracy. AutoSort outperforms existing methods in both simulated and long-term recordings, reducing computational demands by using only 10% of the time and 25% of the memory compared to conventional methods. By combining AutoSort with high-density flexible probes, we track neural dynamics in real-time during motor learning and skill acquisition over 2 months, capturing intrinsic neural manifold drift, stabilization, and post-learning representational drift. AutoSort offers a promising solution for studying long-term neural intrinsic dynamics and enabling real-time BCI decoding.

## Main text

Stable decoding of behavior-dependent neural dynamics from the same cells in the brain is important for brain-computer interfaces (BCIs) as well as for understanding the evolution of neural dynamics during learning, memory, and aging^1^. In addition, tracking the progression of conditions such as epilepsy^2^ or Parkinson’s^3^ disease requires chronic, longitudinal monitoring to fully realize the potential of BCI-based treatment for neurological disorders. Therefore, it is essential to develop stable, high-resolution, fully automated end-to-end systems capable of chronic brain decoding during animal behavior and environmental interactions. However, conventional methods, such as rigid and bulky brain probes, cannot maintain stable recordings from the same cells due to probe drift and immune responses caused by the mechanical mismatch between electrodes and brain tissues. This poses a significant challenge to the stable decoding of neural activities.

Recent advances have tried to address these challenges^4–6^ by providing flexible electrodes with tissue-like mechanical properties and structures^7^. These developments enabled tracking of electrical activity from the same neurons in behaving animals over months to year^8–14^. In addition, the integration of high-density microelectrode arrays with complementary metal-oxide semiconductor (CMOS) multiplexing circuits has created miniaturized brain probes, such as Neuropixels, which can reliably track electrical activity from the same neurons for extended periods, even accounting for any drift during recording^15–17^.

While these hardware innovations are transformative, they introduce substantial software challenges in data management and processing. The ability to maintain stable recordings from the same cells over long periods (months to even years) generates enormous datasets, often ranging from terabytes to petabytes in size, creating complexity in processing, storage, and analysis. A critical bottleneck in this process is converting noisy raw data into identified single-neuron activity, or ‘spikes’ through spike sorting. Despite the potential of spike sorting, enabling a significantly reduced data size due to the sparse nature of spikes^18^, real-time implementation of spike sorting and automatic alignment of spikes from the same neurons over long-time remains a significant challenge, with current approaches relying on *post hoc* processing.

Addressing these software challenges requires a paradigm shift in spike sorting, neuron alignment, and neural activity decoding for long-term stable BCIs. Key requirements include (i) real-time spike sorting for single-cell resolved BCI decoders; (ii) high scalability to handle growing data and complexity; and (iii) accurate and stable alignment of neuronal signals across days to months, facilitating stable behavior-dependent neural manifold decoding for external control or study.

Current spike sorting methods^19,20^ have achieved high accuracy but often rely on human curation and validation for optimal neuron identification, limiting their capacity for automatic and real-time brain decoding. For example, one type of strategy^19^ uses principal component analysis for feature extraction, followed by advanced clustering algorithms such as the ISO-SPLIT clustering algorithm^21^. While effective for single sessions, these strategies require concatenation of recordings and manual curation for longer-term neuron tracking. Although concatenating can apply to long-term recordings^8,13^, it struggles to keep up with the exponentially increasing computational demands as recordings extend to months. For instance, processing data from a 30-channel electrode array over ten days requires over 100GB of peak RAM and more than 2,000 minutes of processing time. Another type of strategy^20,22–26^ focuses on clustering methods like *k*-means or graph-based clustering, integrating template matching and modularity optimization, often enhanced by deep learning. Although this strategy can technically run in real-time by clustering new features into predefined clusters, it struggles to maintain consistent neuron counts across sessions. Recently, some algorithms^25^ have been developed to attempt neuron alignment across sessions for neuroscience studies, but they still lack the consistency in the number of neurons. Additionally, since these algorithms are built on top of existing spike sorting implementations, they have yet to be fully optimized for real-time applications. Thus, no current method fully addresses all three key requirements of real-time, scalability, and consistency simultaneously.

By analyzing stable recordings from flexible electrodes, we identified unique neuronal signal characteristics that could enable real-time stable spike sorting and neuronal signal alignment. Specifically, these recorded data exhibit intrinsic consistency that may help address downstream software challenges. Firstly, neuron spike waveforms demonstrate stable electrophysiological properties, such as shape, amplitude, duration, and slopes, over time. Exploiting these characteristics enhances long-term sorting accuracy, the same as explored by previous template-matching methods^23,25^. In addition, high-density probes allow chronic tracking of waveforms from the same neurons across multiple channels, which intrinsically embeds information about cell location and multi-channel waveform features to improve neuron identification and alignment accuracy. Furthermore, the physical location of neurons relative to high-density electrode arrays remains stable over extended periods. Leveraging this spatial information as a new modality could further enhance the neuron tracking accuracy. Here, we view each of these types of information as distinct modalities, each one providing complementary insights and collectively shaping our understanding of neuron spiking patterns.

We believe that integrating these stable multimodal single-cell resolved features obtained from flexible hardware can significantly improve the accuracy and efficiency of spike sorting and decoding from the same cells during long-term recording. Furthermore, incorporating these multimodal features into deep learning models could enable end-to-end, automated, and *on-the-fly* processing.

To address this need, we introduce AutoSort (Figure 1), a multimodal deep learning pipeline designed for automatic, end-to-end real-time spike sorting and decoding over long-term recordings (Figure 1a). AutoSort is initially trained on manually curated data from the first day of recording (Figure 1b). By tracking the same neuron and aligning them over time using multimodal features we collect from stable signals, AutoSort automatically and accurately performs spike sorting for subsequent days, ensuring consistent neuron tracking throughout months-long recording (Figure 1c,d). We evaluated AutoSort’s performance on various datasets. The result from the simulated electrophysiology datasets with ground truth^27^ demonstrates that AutoSort obtains 99% accuracy. In real datasets, the real-time spike sorting results from AutoSort achieves comparable performance with the state-of-the-art spike sorting. Specifically, compared to offline MountainSort^19^, AutoSort achieved spike detection matches accuracy around 90% and spike classification approximately 95%. By leveraging multimodal deep learning, we enable real-time spike sorting, requiring manual curation only on the initial day, what considerably reduces the time spent by 90% and peak RAM usage by 75% compared to standard methods. This real-time, stable spike sorting, combined with flexible hardware, allowed us to study the evolution of spikes over time. During two-month recording, we used AutoSort to monitor the progression of skill acquisition, capturing three key stages: (i) initial learning dynamics, (ii) stabilization of the acquired skill, and (iii) the representational drift observed after the skill was learned. By applying AutoSort nearly daily, we tracked these changes in real time, allowing us to observe both neural manifold drift and stabilization during motor learning, consistently maintaining accurate neuron tracking throughout the entire process.

**Figure 1.**
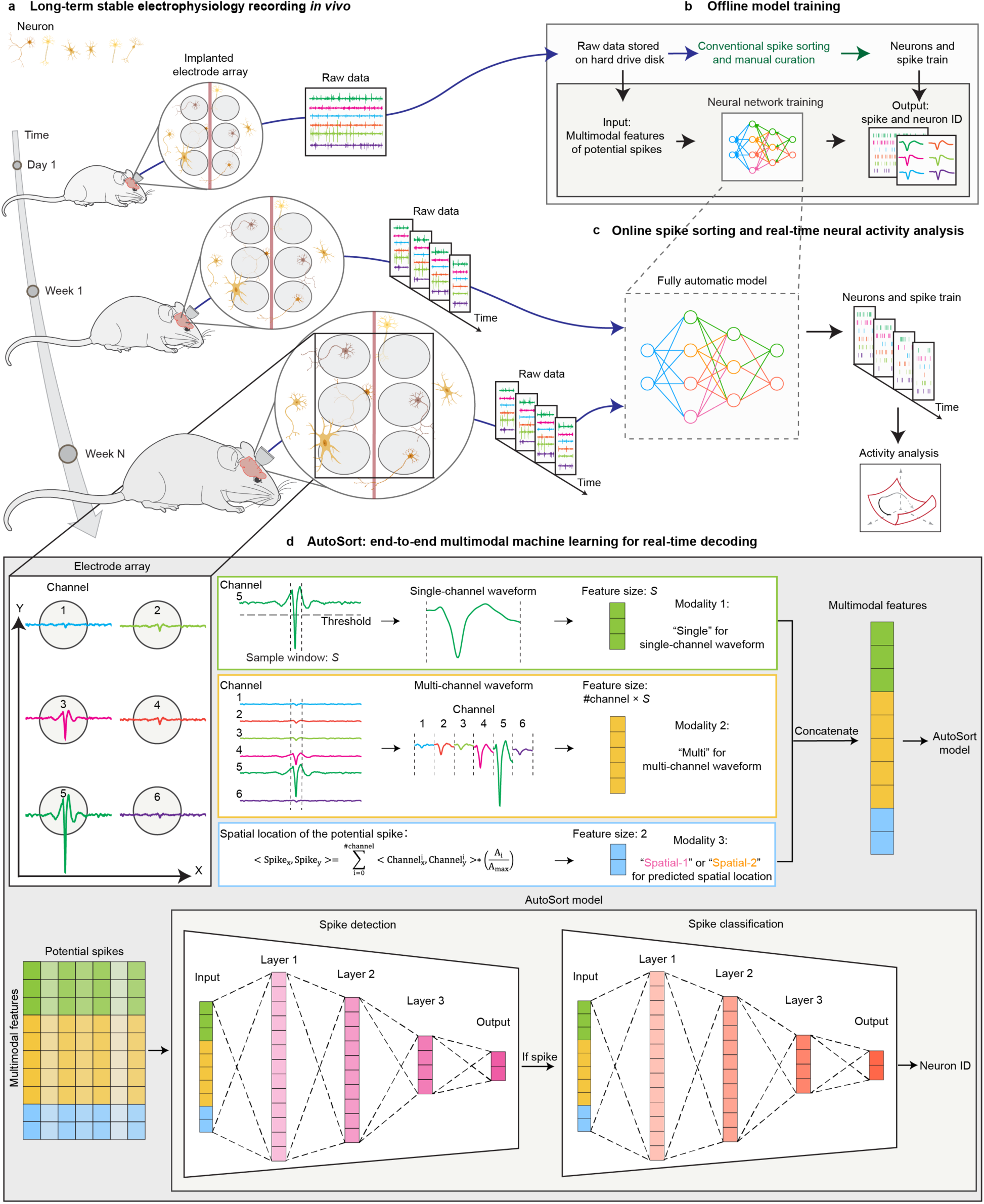
Overview of AutoSort. **a**, Schematics showing the long-term, stable tracking of activities from the same neurons using a tissue-like flexible high-density electrode array during chronic in vivo recordings in behaving animals. **b**, AutoSort model training workflow. Initial stable recording data is processed, curated, and used to train the multimodal model. **c**, End-to-end AutoSort model implementation. The trained AutoSort model enables the automatic and *on-the-fly* decoding of subsequent recording data. **d**, AutoSort model inference. The flexible high-density electrode array detects voltage spikes across multiple channels from a neuron near channel #5. The spike amplitude across different channels varies. Multimodal information is extracted from the multichannel recordings including the single-channel waveform from the channel with the highest amplitude, the multi-channel waveform across all channels, and the estimated spike location based on spike amplitude and channel location. These modalities are concatenated for spike identification and classification.

## Overview of AutoSort

The AutoSort is designed to address two main challenges in long-term stable BCI decoding (Figure 1a). First, it aims to enable scalable, real-time spike sorting while maintaining high precision over extended recordings. Second, it investigates whether integrating multimodal data from high-density probes into spike sorting pipelines can enhance performance using the inherent consistency in stable long-term recordings. Maintaining consistent performance from the initial recordings throughout the later stages without requiring recalibration is crucial for achieving stable decoding.

To address the first challenge, AutoSort trains a deep learning model offline using a small, curated multimodal dataset from initial sessions, such as the recording on the first day, and then applies this model to subsequent recording sessions (Figure 1b,c). AutoSort features a dual-classifier network: the first classifier is responsible for spike detection and noise rejection, while the second one for spike classification. The spike detection classifier determines whether the signal is a true spike or noise. If identified as a true spike, the signal is then classified by the second classifier, which assigns it to a specific neuron. Both classifiers share a common structure, including an adaptive pooling layer, fully connected layers with batch normalization and nonlinear activations, and a final linear layer. The overall training scheme of AutoSort is determined by an overarching loss function that consists of two supervised classification losses optimized simultaneously: one for separating noise and spike clusters during spike detection, and another for separating different neuron clusters during spike classification in latent space. Once trained with initial data, the AutoSort enables scalable, real-time spike sorting of new electrophysiological data from the same animals, allowing for continuous tracking and decoding of neural activity (Figure 1c).

To address the second challenge, we identified key modalities common to all high-density chronic recording probes. In these probes, spikes from a single neuron are often simultaneously detected across multiple nearby channels. For each potential spike, three key modalities can be extracted from the raw data (Figure 1d): (1) the single-channel waveform, defined as the waveform from the channel with the highest amplitude at the moment of the spike; (2) the multi-channel waveform, which concatenates waveforms of the same spike across all channels within the electrode array; and (3) the inferred spatial location, inferred as the coordinates of the neuron, which produces the spike, relative to the electrode array. We demonstrate that combining all three modalities significantly improves accuracy, especially compared to using single modalities alone. Therefore, for any spike that exceeds a specific threshold on a given channel, all three modalities are extracted. For example, the single-channel waveform is captured within a defined spike sample window *S* around the spike event, while the multi-channel waveform combines waveforms from all channels within the same sample window. The spatial location is inferred by incorporating the probe geometry, which provides spatial information for each channel into the multi-channel waveform.

## Optimization of multimodal inputs using simulated data in AutoSort

We optimized the performance of AutoSort by determining the most effective combination of multimodal features as inputs using simulated data with established ground truth. The MEArec simulation tool^27^ was employed to create neuron templates with known ground truths under controlled conditions. The simulation was configured with 50 neurons, with 70% excitatory and 30% inhibitory types, with a minimum spacing of 25 micrometers between neurons. The system was set to simulate a zero-noise environment with an overlap threshold at 0.8. This threshold decides when two templates are considered overlapping. For example, with the setting of 0.8, two templates overlap if Template B’s signal is at least 80% as strong as Template A’s strongest signal on the same electrode. These neuron templates are used to generate spike trains, which facilitated the creation of simulated extracellular recordings (Extended Data Figure 1a).

We applied AutoSort models to these simulated recordings to assess their performance (Extended Data Figure 1b). These recordings intrinsically define single-neuron waveforms and multichannel waveforms. We tested the accuracy of the AutoSort using four distinct input modalities: (1) single-channel waveform (*Single*), (2) multi-channel waveform (*Multi*), (3) predicted spatial location using Scheme 1 (*Spatial-1*), and (4) predicted spatial location using Scheme 2 (*Spatial-2*) (Figure 2). We observed that in a stimulated, zero-noise environment, the ideal multi-channel waveforms of a potential spike showed distinguishable spike waveforms only on channels near the actual location of the neuron, with amplitude decreasing proportionally to the distance between the channel and the neuron. For the spatial information, we explored two distinct schemes to infer the coordinates from these waveforms, as illustrated in Extended Data Figure 2a-b. Under these ideal conditions, *Scheme 1*, which directly weighs the position of each channel by the waveform amplitude to estimate the spatial location of potential spikes, provided a reasonable prediction of neuron location (Methods). However, in more realistic, noisy data conditions, noisy amplitudes could often make Scheme 1 (*Spatial-1*) produce inaccurate predictions. To address this limitation, Scheme 2 (*Spatial-2*) was designed to focus on local waveform characteristics, estimating the spike’s origin within a specified radius based on the amplitude and spatial distribution of the signal peaks. This approach refines the prediction by focusing solely on the nearest channels around the channel where the maximum amplitude is identified (Methods). By considering the localized impact of the neuron’s activity, the method aims to enhance prediction accuracy especially in noisy environments (Extended Data Figure 2a-b).

**Figure 2.**
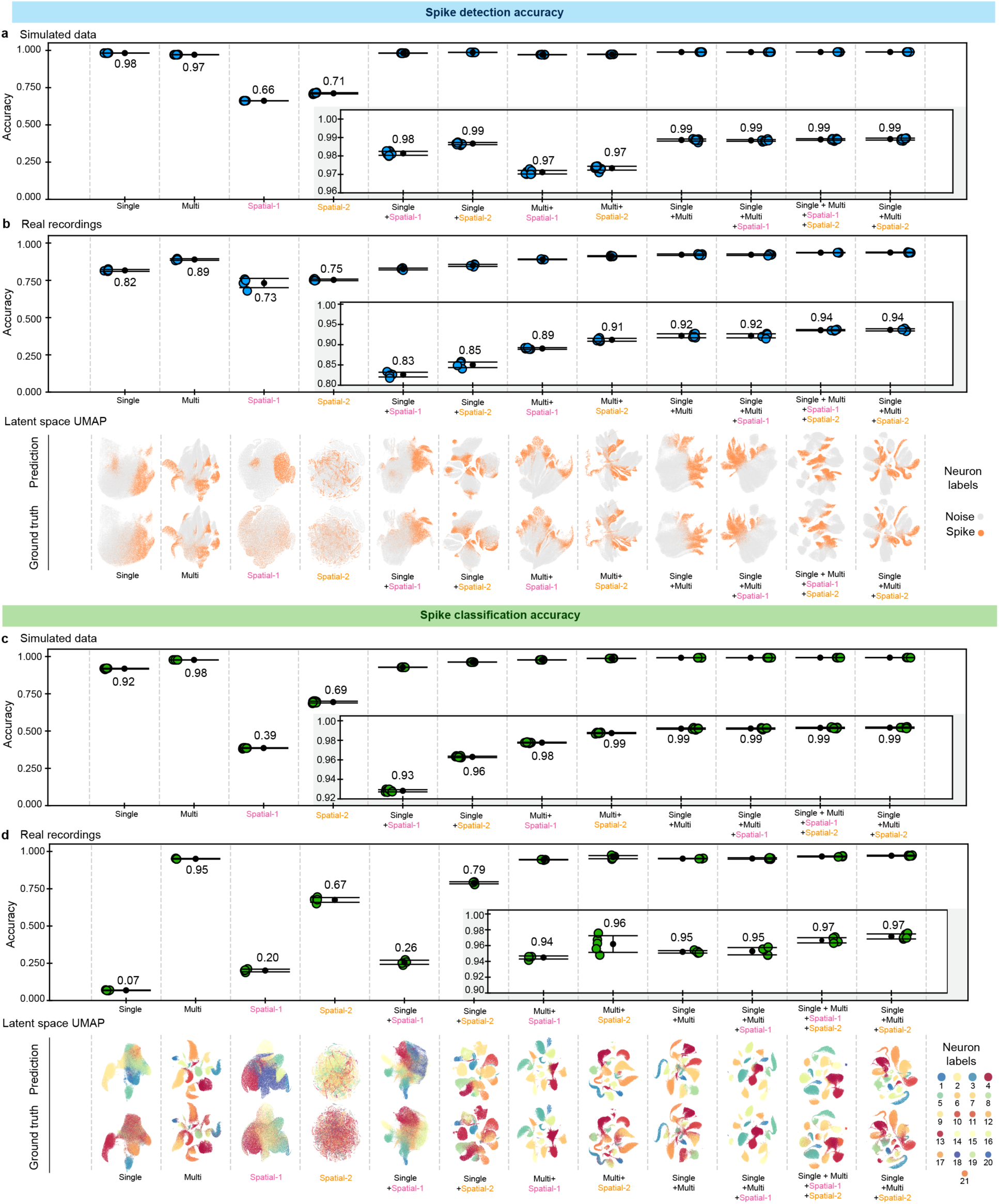
Evaluation of AutoSort performance with different combinations of input modalities on simulated and real electrophysiological data. **a**, Spike detection accuracy on simulated electrophysiological data across different seeds (n = 10) of AutoSort with different combinations of input modalities. **b,** Spike detection accuracy on real electrophysiological recording data across different seeds (n = 5) of AutoSort using the same input modality combination as in (**a**). The bottom panel shows corresponding latent space UMAP plots of spike detection under the same input modality combinations as in (**a)**. **c,** Spike classification accuracy on simulated electrophysiological data, as in (**a**). **d,** Spike classification accuracy on real electrophysiological data, as in **(b)**. All models for the real electrophysiology dataset were trained using Day 6 (after 21 days post implantation) as the training data point, and Day 38 (after 21 days post implantation) as the representative test data points. Colored dots represent individual accuracy values for each AutoSort model, and the black dot represents the average. Error bars represent standard deviations (SD).

Results showed that the *Single* or *Multi* modalities contained more information than the *Spatial-1* and *Spatial-2* modalities. Notably, the *Single* modality was more effective for spike detection but less informative for spike classification compared to the *Multi* modality. This suggests that the signal distribution we obtain through the multi-channel waveforms provide more information for neuron identification than the single-channel waveforms alone. Additionally, while using the *Spatial-1* and *Spatial-2* modalities alone has relatively low accuracy for spike sorting, the *Spatial-2* modality showed improved accuracy over the *Spatial-1* modality.

Next, we examined whether combining two and more modalities as input would be beneficial (Figure 2a, b). We find that combinations including *Single* + *Spatial-1* or *Spatial-2*, and *Multi* + *Spatial-1* or *Spatial-2* resulted in higher accuracy than single modalities alone. Additionally, comparisons between combinations including *Spatial-1* versus *Spatial-2* consistently showed that combinations with *Spatial-2* resulted in improved accuracy. Finally, we explored accuracies with multiple modalities. Adding *Spatial-1* to *Single* + *Multi* provided minimal improvement in accuracy, but when *Spatial-2* was added, the accuracy was more increased. In conclusion, the results show that the information in each modality was not fully overlapped so that combination of different modality can improve the spike sorting accuracy. The combination of *Single* + *Multi + Spatial-2* produced the highest and stable results.

Additionally, we investigated how different experimental variables affect spike sorting accuracy in AutoSort. Using the simulation tool MEArec, we mimicked various experimental settings by adjusting experimentally relevant parameters, including the number of neurons (from 20 to 100, Setting 2), the threshold to consider two neurons spatially overlapping (from 1.0 to 0.0, Setting 3), the minimum distance between neurons (from 500 to 25 μm, Setting 4), the variance of additive Gaussian noise (from 0 to 40 μV, Setting 5), and cell types (four excitatory neuron types and four inhibitory neuron types, Setting 6; Extended Data Figure 3a). The results showed that as the number of neurons increased, the accuracy of spike sorting slightly decreased (Extended Data Figure 3b). This aligned with our expectation, as increasing the number of neurons recorded per channel will make it more challenging to accurately identify and classify spikes into the correct neurons. Also, as the minimum distance between neurons increased, we observed a decrease in accuracy (Extended Data Figure 3e). In contrast, variations in the overlap threshold, the variance of additive Gaussian noise, and cell types (Extended Data Figure 3c,e,f) did not impact the performance. Together, AutoSort demonstrated robustness maintaining the accuracy over 94% in a variety of settings.

## Application of AutoSort for months-long stable in vivo electrophysiology

We further examined the effectiveness of AutoSort by applying it to long-term, stable in vivo electrophysiological recordings in behaving mouse brains. Using our previously reported mesh electronics with tetrode-like electrode arrays^6^ (Extended Data Figure 4), we tracked single-unit action potentials from the same neurons over time in mice. We first trained AutoSort on recording data from an early first time point (Day 6, after 21 days of flexible electrode array implantation in Mouse #1). The training data was processed by MountainSort followed by human curation. We then applied the trained AutoSort model to later-stage recordings (Day 11, 16, 27, 38, 50, after 21 days of flexible electrode array implantation in Mouse #1) for automatic spike sorting. To test its performance, we further used MountainSort and human curated spike sorting results as ground truth.

We evaluated the accuracy of the trained AutoSort model in the second-time point recording through the similar tests on the simulation data (Figure 2a, c) focusing on how different combinations of input features affect the accuracy (Figure 2b, d). Using the data from Day 38 as an example, first, as the *Single* or *Multi* modality alone is less effective, showing that single modality cannot achieve high accuracy in real recording data. (Figure 2d). Second, combining multiple modalities produced consistently better results. The combination of *Single* + *Multi* + *Spatial-2* produced the best results, having a matching accuracy with MountainSort of 94% in spike detection and 97% in spike classification. Finally, to further illustrate the performance of different modality combinations, we plotted the Uniform Manifold Approximation and Projection (UMAP)^28^ of latent space in the last layer of spike detection or spike classification classifier trained with different input modality (Figure 2). The latent space from the model using a combination of modalities revealed improved separability between noise and spikes as well as between spikes from different neurons.

We used the flexible mesh electronics to stably track the activities from the same neurons for 50 days. We further evaluated the performance on additional time points during this long-term recording and confirmed that combining multimodal information consistently improved accuracy (Figure 3). The combination of multimodal inputs achieved higher accuracy, with the *Single* + *Multi* + *Spatial-2* combination standing out as the highest, reaching over 90% in spike detection and well over 92% in spike classification, in both simulated and real electrophysiological data. Similar results were obtained from the long-term recordings of Mouse #2 (Extended Data Figure 5a,b), where accuracies consistently exceeded 90% and 95%. As a result, this multimodal combination approach was selected for subsequent simulation data analyses.

**Figure 3.**
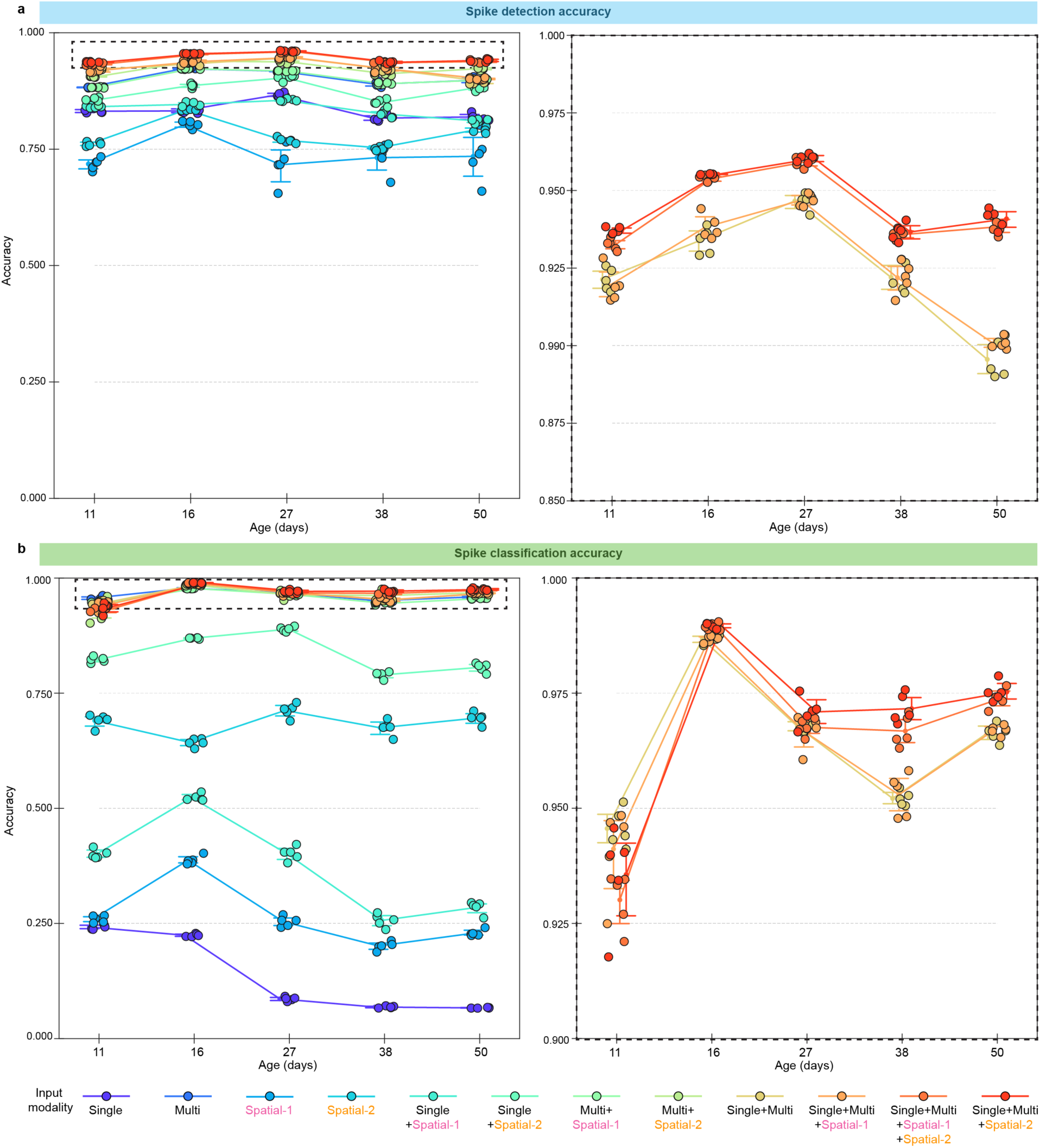
Evaluation of AutoSort performance for chronic electrophysiological recording data analysis. **a**, Spike detection accuracy of AutoSort with different combinations of input modalities on representative chronic electrophysiological recording data. On the right, the highest-performing spike detection models are zoomed in for comparison. **b**, Same as in (**a**) but for the spike classification accuracy. To train all the models, we used day 6 (after 21 days post implantation) as the training data point and tested on days 11, 16, 27, 38, and 50 (after 21 days post implantation) as representative data points. For all the strip plots, dots and line plots represent the mean accuracy values across the different seeds (n = 5) of the models for each test data point, and error bars indicate the standard deviation.

We also explored whether AutoSort could be effectively applied to process streaming recording data in real-time while maintaining high accuracy with multimodal feature inputs. To test this, we extended AutoSort to an online workflow for real-time spike sorting (Figure 4a). We assessed the performance by comparing the matching accuracy between AutoSort and MountainSort, both every second and as a 10-second average, in both online and offline configurations (Figure 4b,e). Overall, we observed that online AutoSort performed comparably to offline AutoSort in spike classification, achieving over 90% accuracy, but worse in spike detection. However, in spike detection, although the sparsity of the spikes initially led to some accuracy drops with occasional outliers, these were minimized when accuracy calculations were extended over 10-second intervals. Also, we can see how the latent space UMAP of online AutoSort, in both tasks, results gradually shaped to ground truth as the recording time increased (Figure 4d,g). Next, we examined the online accuracy in the long term (Figure 4c,f) and confirmed that the accuracies are on average over 80% accurate in spike detection, independently of the day of testing. On the other hand, for spike classification, the performance was very closely matched with the offline version, having performance around 90%. Finally, we also compared online AutoSort results with offline AutoSort in the long-term recordings of Mouse #2. In this second dataset, online AutoSort also consistently maintained high accuracy across different testing days, achieving over 70% accuracy in spike detection and well over 90% in spike classification. (Extended Data Figure 5).

**Figure 4.**
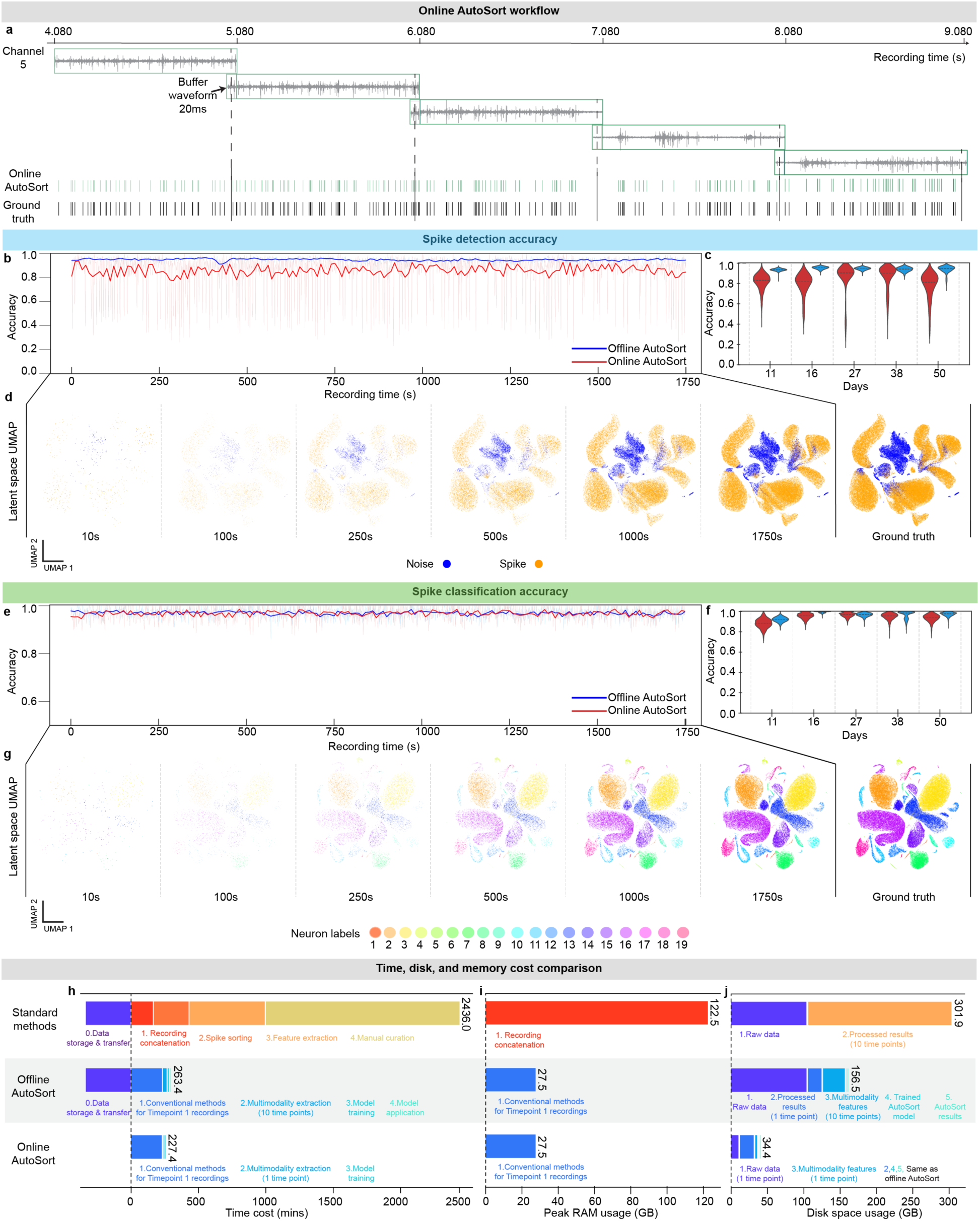
AutoSort-enabled automatic online spike sorting for long-term stable electrophysiological recording in behaving animals. **a,** Representative workflow showing the AutoSort-enabled online spike sorting from a five-second in vivo recording on one channel of a mouse, seven weeks post-implantation. **b-d,** Comparison of spike detection accuracy between online (red) and offline AutoSort (blue). **b,** Spike detection accuracy for each second of recording on Day 27 (after 21 days post implantation) in lighter blue and red shows the 1-second accuracy over time, and in darker red and blue, the 10-second averages of online and offline AutoSort, respectively. **c,** Violin plots of the spike detection accuracy comparison between online and offline AutoSort on the representative chronic recording data in the representative animal from Days 11, 16, 27, 38, and 50 (after 21 days post implantation). **d,** Latent space UMAP of online AutoSort spike detection results as recording time increases. **e-g**, Same as in (**b-d**) but with the accuracy of spike classification. **e**, Same as in (**b**) but with spike classification accuracy for each second of the same recording. **f**, Same as in (**c**) for the spike classification accuracy **g,** Same as in (**d**) but with the spike classification accuracy. **h-j**, Computational Resources comparison between offline AutoSort, online AutoSort, and MountainSort (Conventional Methods). Benchmarking calculations and estimations using 1 day for training and 9 different days for data points, highlighting the savings with increased data for spike sorting. **i,** Cost of running time needed for the spike sorting process. **i,** Cost of memory usage for the spike sorting process. **j**, Storage space needed for the spike sorting results. For all violin plots, the line inside the violin plots represents the median, providing an overview of the data accuracy distribution.

In addition to maintaining high accuracy, we compared the time, memory, and storage cost of online and offline AutoSort with currently used methods^19,29^ (Figure 4h-j). When comparing offline AutoSort to conventional methods, we observed a considerable saving in time, peak memory, and ROM cost because offline AutoSort only requires processing on Time point #1 recording sessions with conventional methods as training data to train the model. In terms of time cost (Figure 4h), AutoSort could largely bypass the time-consuming procedure associated with concatenating multiple time-point recordings, spike sorting, feature extraction, and manual curation, which are typically required by conventional methods. In addition, because the computation-intensive task of recording data concatenation is not necessary, peak RAM usage is substantially reduced from 122.5 GB to 27.5 GB (Figure 4i).

However, while offline AutoSort greatly improved time, and memory cost, it still required raw recording data to be stored. After the recording session ends, this data must be transferred to a workstation for processing, leading to additional time cost and increased disk space usage. To address these issues, online AutoSort further optimizes data storage, transfer time (Figure 4h) and disk space by processing raw data during the recording process. Instead of raw recording data, only sorted neurons and spike train results are processed and saved, reducing disk space usage from 301.9 GB to 34.4 GB (Figure 4j).

## AutoSort enhances decoding of months-long single-unit action potentials during learning of a motor task

Online AutoSort enables real-time neural population dynamics analysis, essential for immediate feedback in dynamic experimental settings. The real-time capability not only reduces the data management load by processing data *on-the-fly*, enhancing efficiency and scalability, but also allows ongoing model adjustments based on live data to maintain decoding accuracy. Moreover, real-time application in animal learning studies enables real-time tracking and analysis of how neural patterns evolve with new experiences, thus offering insights into learning synaptic plasticity and memory mechanisms^30–31^.

To explore the potential of online AutoSort in tracking the neural mechanisms of motor learning, we processed electrophysiological recordings during mouse motor learning. We implanted mesh electronics with tetrode-like electrode arrays into the mouse motor cortex. After 21 days of implantation, the animal was trained to reach a joystick to obtain a water reward (Extended Data Figure 6). We recorded dynamics of single neurons across nearly daily sessions and tracked the mouse’s learning and proficiency in the task from Day 1 to Day 50. First, four days of recording data was processed, manually curated, and used to train AutoSort, ensuring all the recorded neurons have been included in the training. Once trained, we utilized it to analyze the neuron activity as the mouse began learning and eventually mastered the motor reaching task. For a comprehensive evaluation, AutoSort was used in real-time over subsequent days, automating the spike sorting process. We compared the performance of AutoSort in this online application with its offline use and an online spike detection-only method without sorting, assessing the efficacy of each in identifying specific neuron activity.

Using online AutoSort, we obtained sorted neuron spike trains (Figure 5a). We then assessed the ability to capture low-dimensional neural representations^32^ by comparing the initial learning stage trajectories with those after the mouse achieved proficiency (Figure 5b). During learning stages from Day 1 to Day 31, all models showed constantly evolved neural trajectories, particularly the offline AutoSort, which is expected to be the most accurate. We observe that these trajectories after Day 31 became more consistent as animal training continued. Pearson correlation of daily neural activity with the average for each phase to differentiate between the learning and after learning periods showed that the offline AutoSort methods exhibited higher correlations across all principal components (PCs) during proficient days (Figure 5b-c).

**Figure 5.**
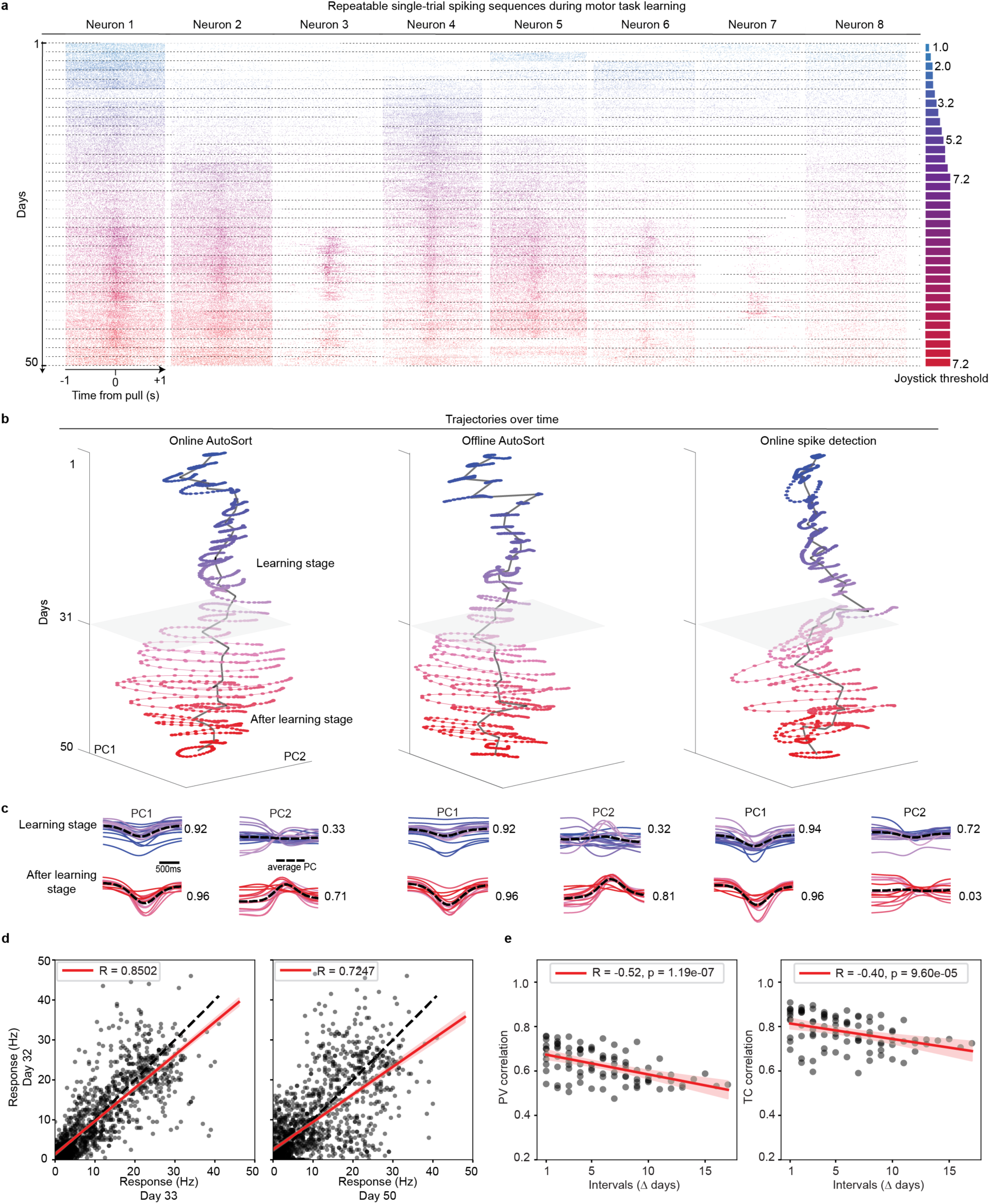
AutoSort enables online behavior decoding during motor learning and skill acquisition from the same cells. **a,** Visualization of representative individual neuron spike trains during the learning process, color-coded to show the transition from the learning phase (blue) to the after learning phase (red). Each day displays 50 representative trials of spikes during the motor task recorded. To the right, the joystick threshold level per day of the animal experiment is also shown. **b,** Trajectory analysis of neural activity within a two-second interval, averaged over trials each day from day 1 to 50 (after 21 post implantation) throughout both the learning and after learning phase. From left to right, the trajectory plots include the online, and the offline AutoSort models, and an online spike detection model, where each spike originates from one channel without neuron sorting. Each trajectory is a low-dimensional representation of daily trial spikes, with colors consistent with (**a**). The gray line represents the average coordinate of the manifold for each day. **c,** Decoupled PC1 and PC2 for each sorting method, with a dashed line representing the average PC trajectory per each learning stage model. **d,** Correlation between neurons’ firing rates obtained through online AutoSort at the first day after learning (y-axis) and those at later time (x-axis). Each dot represents one neuron’s mean firing rates at the two different times. The black dashed line is the unity line, and the red line with shade is the regression line plotted with 95% confidence interval (CI). **e,** The correlations of PVs, and TCV between all pairs of recording sessions, using online AutoSort during the after learning stage, plotted against the corresponding time intervals. Each dot represents one correlation at two different times. The straight lines with shades are the regression lines with 95% CI of the linear regression.

Moreover, among all methods - offline, online with sorting, and online without sorting - PC1 showed the greatest alignment after learning, with Pearson correlations of 0.92, 0.92, and 0.94 during learning compared to 0.96, 0.96, and 0.96 after learning. For PC2, the offline and online AutoSort methods demonstrated similar Pearson correlation changes before and after learning, with consistent behaviors observed (0.32 and 0.33 during learning vs. 0.81 and 0.71 after learning). In contrast, the online spike detection-only method without sorting did not show the same behavior after learning (0.72 during learning and 0.03 after learning). As the mouse proficiency increased, PC2 showed higher correlations in the offline and online methods during the post-learning phase, while the method capturing spikes without sorting did not (Figure 5c). Our results show that by sorting the neurons, either online or offline, we can achieve more stable trajectories in late stages, once the animal has been more exposed to pulling the joystick. These findings suggest that neuron sorting might be beneficial for achieving consistent neural representations during motor tasks, in line with previous literature^33^ studying the impact of spike sorting.

With the same sorted spikes obtained with online AutoSort, we computed the stability of motor representation in the motor cortex over time, focusing on the post-learning period to minimize variability from the learning process (Figure 5b-c). Neural representations of motor intentions were highly correlated between consecutive days, such as Day 32 (the first day after the learning stage) and Day 33 (R = 0.85), indicating stable motor representations (Figure 5d). However, as the interval between recordings increased (e.g., between Day 32 and Day 50), the correlation decreased (R = 0.72), suggesting representational drift could also exist in the motor cortex over time. This drift was quantitatively measured by calculating the correlation of population vectors (PV) and tuning curves (TC) against the intervals between sessions. The negative correlations observed for both PV (R = -0.52, p = 1.19 × 10^-7^) and TC (R = -0.40, p = 9.60× 10^-5^) indicate a gradual reduction in representational similarity as the interval between sessions increases, highlighting the dynamic nature of motor cortex representations and the importance of spike sorting for capturing stable representations (Figure 5e).

## Discussion

We introduced AutoSort, a multimodal deep learning-based algorithm designed for automatic, scalable, and real-time spike sorting and neuron tracking in chronic stable electrophysiological recordings. AutoSort enables end-to-end tracking by continuously aligning the same neurons over time through multimodal features derived from high-density recordings, ensuring consistent performance across sessions. We demonstrated that multimodal combinations of single-channel waveforms, multi-channel waveform distributions, and spatial consistency of neuron locations achieved higher, more robust and more consistent performance in long-term spike sorting compared to single modality inputs. This multimodal deep learning approach also allows for real-time inference, notably reducing the manual effort and computational load required by traditional methods. Moreover, this stable real-time decoding, combined with flexible, high-density hardware, enables the long-term study of neural dynamics. For example, we demonstrated its application in motor learning and skill acquisition, where after the stabilization of neural manifolds, we observed representational drift post-learning in the motor cortex, as observed in other brain regions. This advancement facilitates more scalable and sustainable studies of neural activity, supporting online and on-chip data processing and compression.

While AutoSort can manage the vast data volumes produced by long-term neural recordings, several directions for improvement remain. The initial need for manual curation introduces potential inaccuracies that can propagate throughout the entirety of the data analysis, potentially leading to errors in neural activity interpretation. Additionally, the current framework demands the recording system to be connected to a computer for real-time data processing and sorting, which limits the feasibility of studies on naturalistic environments, sleep patterns, or simultaneous recordings from multiple animals. To address these issues, integrating AutoSort into specialized hardware platforms could offer a solution. Neuromorphic frameworks and technologies^34–36^, which mimic the neural structures and processing methods, hold promise for further improving efficiency while maintaining the accuracy in real-time long-term BCI systems.

Looking ahead, integrating out-of-distribution detection^37^ techniques into AutoSort could enhance its ability to manage new and unexpected neuron appearances by adapting to changes and correcting errors. These techniques would facilitate the detection of new neurons over time, particularly when training data is limited compared to the extensive testing period, such as training on one day of data while testing over several months or years. This approach, utilizing semi-supervised learning and iterative self-training, could improve the system’s reliability and long-term performance. However, despite its potential, AutoSort has several unsolved limitations. The system performs optimally when the data is stable, and the training dataset is well curated. Yet, if the data is unstable, such as when there is noticeable drift of signals on the neural probes, the system’s accuracy may be compromised. Integrating out-of-distribution detection and adaptive learning mechanisms could address these issues by continuously monitoring and adjusting to changes in neural activity, thus enhancing the system’s resilience and accuracy in long-term recordings. Additionally, in recordings with pronounced probe drift, such as those from NeuroPixels, incorporating algorithms to account for signal drift^25,38–39^ is crucial for enhancing system stability before applying AutoSort. Nevertheless, maintaining highly stable recordings with consistent neuron counts over months and years remains essential. If consistency cannot be ensured, missing data may need to be addressed through simulations or spike-based foundational models^40^ to ensure that the neural manifolds are consistently derived from the same neuron population.

## Methods

### AutoSort models

For the spike detection and spike sorting task, two classification models (M*_noise_* and M*_cls_*) are used for the spike detection and spike classification task respectively. The input spike is denoted as *X* ∈ R*^n^*^×*p*^, where *n* is the data number and *p* is the dimension. *X* is concatenated with other low dimensional features such as spatial information to generate *X*′ ∈ R*^n^*^×*d*2^ as the input of M*_noise_* and M*_cls_*. These two models are trained simultaneously with Binary Cross-Entropy (BCE) Loss for the spike detection and spike classification tasks. The details of the algorithm are as follows.

## Classifier architecture

Both the spike detection and classification models have the same architecture, differing only in the number of output classes. The spike detection model has two output classes, corresponding to whether the spike is noise or not, and the spike classification model has a number of output classes equal to the number of neurons being measured. Both classification models are structured as feed-forward neural nets with five layers. The first layer is an adaptive pooling layer, and the next three are fully connected layers with batch normalization applied before a ReLU activation function. The final layer is a fully connected linear layer. Both classifiers are trained with BCE loss, so applying a sigmoid function to the output logits directly produces the classification result.

## End-to-end model inference and training

To train the AutoSort model, initially, an input feature vector is created for each potential spike detected. This vector comprises the recorded waveform, a combination of multiple waveforms, and the spike’s estimated spatial coordinates, as depicted in Figure 1e. The classifiers within the model are designed to assess these inputs, determining the authenticity of each spike and identifying the originating neuron sources for genuine spikes. The training strategy incorporates a dual-component loss function: one part focusing on binary cross-entropy (BCE) loss for filtering out noise (noise rejection BCE loss), and the other on BCE loss for accurately classifying actual spikes (spike classification BCE loss), with the latter applied solely to validated spikes. The Adam optimizer is employed to fine-tune the model, utilizing back-propagation based on the cumulative BCE losses. During this phase, potential spikes are identified, and three distinct types of information are gathered for each. These spikes are labeled as true (for those associated with sorted neurons) or false. Additionally, each true spike is tagged with a neuron ID. This labeled dataset enables the AutoSort model to be trained to perform two key functions: rejecting noise-associated spikes and classifying authentic spikes into their respective neuronal groups. This learning process equips the model to differentiate and exclude spikes caused by noise while accurately assigning genuine spikes to the correct neuron categories based on the provided information modalities.

To perform inference, once the AutoSort model is fully trained, it is applied to new, unseen data for inference purposes. During this phase, the model leverages its training to instantly identify and sort spikes, distinguishing between noise and real neuronal activity. The outcome is a direct and precise classification of spikes into sorted neurons, facilitating the analysis of single-neuron activity during motor tasks. This two-step approach ensures that, after the initial training, the AutoSort model can be utilized to process new datasets, providing sorted spikes for further examination without the need for manual intervention.

## Multimodal features

For training as well as inference purposes, a vector of input features is generated for each potential spike. Potential spikes are detected using a thresholding approach, where the threshold is set to three times the standard deviation of the preprocessed and filtered signal. When this threshold is exceeded, the generated inputs from this spike detection include a single measured waveform, concatenated multiple waveforms, and the predicted spike coordinates (Figure 1d).

Specifically, the waveform feature is constructed by aggregating 1ms preceding and 2ms subsequent data points, creating a 3-ms-point waveform across all recording channels. Additionally, spatial predictions are computed using multi-channel waveform data and electrode locations (Spike*_X_,* Spike*_Y_*), based on the probe’s geometry of choosing (Extended Data Figure 2). This results in each feature vector having a dimensionality of 30 samples × (number of channels + 1) + (dimension of the spatial coordinates) (Extended Data Figure 4).

## MEArec simulations

The MEArec^27^ package is utilized to evaluate the spike sorting capabilities of the AutoSort model under a variety of simulated conditions. MEArec is a tool combining machine learning with biophysical simulations that enables the creation of highly realistic brain activity simulations. These simulations serve as a benchmark for training and assessing the AutoSort model across diverse settings and specifications.

To generate these simulations, MEArec first constructs templates based on various cell models randomly positioned around the probe. These templates then facilitate the production of simulated recordings through a multi-step process. Initially, spike trains are generated following user-defined settings. Subsequently, these trains are convolved with selected templates using amplitude modulation to introduce physiological variability. The final step involves the addition of noise, which can be filtered according to specific requirements. These detailed simulations (Extended Data Figure 1a) enable the comprehensive training and testing of AutoSort models under controlled yet varied conditions.

The evaluation process involved adjusting several key parameters: the number of neurons, the cell model types (including ‘STPC’, ‘TTPC1’, ‘TTPC2’, and ‘UTPC’), the electrode distance (ranging from 10 to 2,500 micrometers, with the distance parameter controlling the electrode separation), and the modality of the input (single or multiple). These simulations are conducted on a 32-channel setup, with the number of neurons varying from 20 to 100. Through this methodical approach, the impact of each variable on the AutoSort model’s performance is dissected and quantified, paving the way for optimized configurations in real-world applications.

## Data acquisition and ground truth spike sorting labeling for the long-term recordings

Daily neural activity recordings (N=2) from week 1 to week 9 (after waiting 21 days post-implantation) are conducted using the Intan Technologies RHD2132 amplifier chip, coupled with a custom printed circuit board (PCB) for connectivity. Recordings are taken at a 10 kHz sampling rate, using tetrode-like mesh electronics for approximately 20-minute periods. Data preprocessing is performed using the SpikeInterface^29^ framework, which applied bandpass filtering to remove noise and irrelevant frequencies, followed by common average referencing using a global reference to reduce artifacts. The recordings are concatenated across sessions for consistency. For the first days, spike sorting is then carried out with the Mountainsort4^19^ algorithm, implemented through SpikeInterface, where spikes are detected and clustered from the preprocessed signals. Units are manually curated using SpikeInterface’s *NpzSortingExtractor*. Waveforms are extracted using SpikeInterface’s waveform extraction module with a 1 ms pre- and 2 ms post-spike window. These curated units, validated by metrics such as interspike interval (ISI) variability, autocorrelograms, electrode spatial information, and waveform amplitude consistency, are considered ground truth for AutoSort evaluation. Long-term stability of the single-unit recordings is confirmed based on spike waveform consistency, and estimated neuron positions across sessions.

## Real-time processing

For real-time acquisition, a similar approach is followed. The RHD2132 electrophysiology amplifier chip recorded signals from 30 channels at a 10 kHz sampling rate. Data is saved in real-time using the ‘One Folder per Signal Type’ format, separating global information into a standard Intan RHD header file and saving waveform data for each channel in separate raw data files. A custom Python script reads and processes this data in real-time. The script reads samples of timestamp data, converting timestamps from bytes to seconds, and stores them as keys in a dictionary for corresponding channel data. For each channel, the script scales and offsets the data from bytes to millivolts and adds it to the dictionary. This process continues until all data is read and processed. If the recording exceeds the buffer (3 minutes), the script cleans the memory of the lists, storing timestamps and data values to prevent excessive memory usage and potential crashes.

## Motor behavioral animal training

Mice are trained in our motor skill learning assay (Figure 5a) across nearly two months of sessions (Day 1-17, 25-30, 32-35, 37-45, 49-50 after 21 post implantation days, following initiation of training in both Mouse #1 and #2). The force required to press the joystick is constant, and the deflection amplitude threshold required for registering a ‘trial’ is increasing from 1 mm to 7.2 mm, in the span of the whole experiment, although after 1 month it is constant at the maximum threshold value. Animals are either transferred to the training boxes for 20 to 30 minutes training sessions nearly 5 times a week or kept in the animal facilities. The joystick-press sequences are shaped by rewarding water-restricted mice with water for approximations to the desired sequence (1,500 ms apart for all mice). The predefined and automated training protocol is implemented in custom software (C#) and hardware.

## Behavior animal task data analysis and neuron-level representational drift

After spike sorting with AutoSort to isolate single spikes from neurons, these spikes are correlated with 2-second-centered joystick movements to compile spike trains for the sorted neurons. A Hebbian/anti-Hebbian neural network^32^ is used where an online dimensionality reduction and dynamic feature extraction conduct a single-layer neural network that linearly maps neural activity to principal component scores. The artificial network, with lateral connectivity matrix *M* and feedforward matrix *W*, projects input data vector *x* from *R^D^* space to a lower-dimensional space *R^k^*. The weight matrix *W* updates according to the rule *W*← *W* + η(*yx*^T^ − *W*), where η is the learning rate. Following this, data is smoothed using a Gaussian kernel 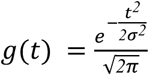, with the *y*_out_ derived by convolving reduced-dimensionality data *y* with *g*(*t*), applying temporal smoothing to the dimensionality-reduced neural activity data.

After extracting the principal components (PCs) for each trial, a daily global PC trajectory is derived by averaging these components for each respective day. The mean PCs for the two distinct phases of the learning process - the learning phase and the post-learning phase - are then determined. The correlation between the daily global PC trajectory and the average PCs for each phase are quantified by computing the Pearson correlation coefficient for every component with its learning phase average PC.

Spike trains are analyzed for neuron-level representational drift by utilizing established methods^30–31^, including population vectors (PVs) and tuning curve vectors (TCVs). Mean firing rates for each neuron are computed for each 0.5-second time bin within the 2-second trial by counting spikes and dividing by the bin duration. Each PV represents the firing rates of neurons in response to a single 0.5-second time bin, while each TCV captures the average firing rates of individual neurons across all time bins during the trial. Pearson correlation is then computed for both types of vectors across different pairs of sessions.

## Data availability

Data used in the study is uploaded to Figshare and will be public with the manuscript publication.

## Code availability

The AutoSort software is available at http://github.com/LiuLab-Bioelectronics-Harvard/AutoSort. Code used in the study is uploaded to Figshare at https://figshare.com/s/8229c4a53e41d5a231f2 and will be public with the manuscript publication.

All data analysis and visualization in this study were implemented based on Python 3.10.13 and MATLAB_r2023a. The following packages and software were used: SpikeInterface 0.101.1 (https://github.com/SpikeInterface), Jupyter Notebook 7.2.2, TensorFlow 2.15.0, Seaborn 0.13.2, Pandas 2.2.1, SciPy 1.14.1, NumPy 1.26.4, Scikit-learn 1.4.1.post1, and MountainSort 4 (https://github.com/flatironinstitute/mountainsort).

## Acknowledgements

Y.H. acknowledges the James Mills Peirce Fellowship from the Graduate School of Arts and Sciences of Harvard University. A.M-L. acknowledges the support from the RCC-Fellowship of Harvard University.

## Author contributions

J.L., and Y.H., conceived the idea. Y.H. developed the method and performed mice implantation and motor behavior training. Y.H. and A.M-L. designed computational data analyses, prepared figures, and drafted the manuscript. H.S. contributed critical discussions and input on the figures. R.L. fabricated the flexible mesh electrodes. Y. H., A.M-L., and J.L. revised the manuscript. J.L. supervised the study.

---

**Extended Data Figure 1.**
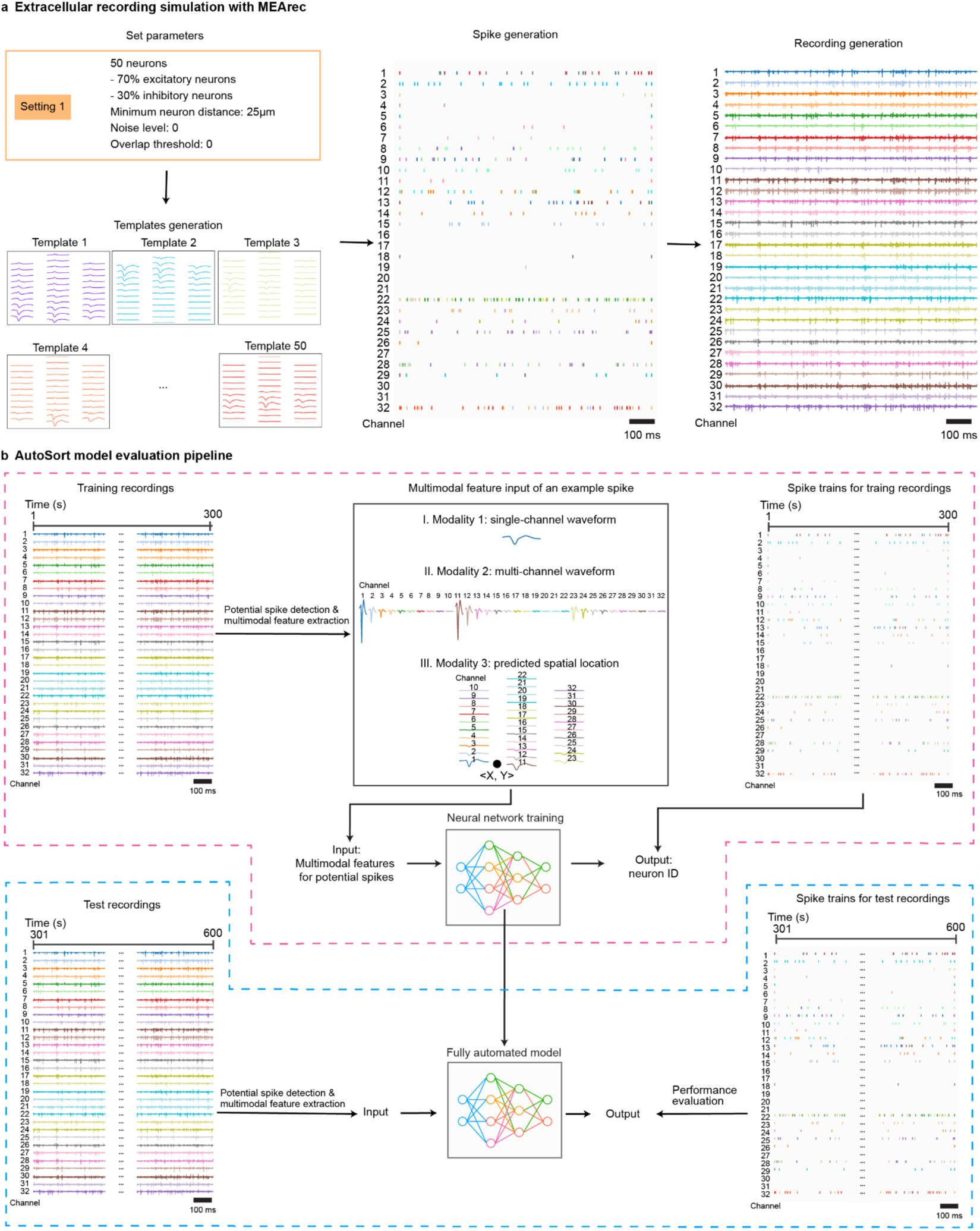
Simulated extracellular recording data end-to-end workflow. **a,** Extracellular recording simulation workflow with MEArec^27^. With user-defined parameters, MEArec generate templates, which is used to generate spike trains and electrophysiological traces. **b,** AutoSort model pipeline on simulation data. The 10-min simulation data is separated into training (1-300s) and test (301-600s) recordings. Training data and its corresponding ground truth spike trains are used to train a model. Then test data and corresponding ground truth spike trains are used to evaluate performance of the model on MEArec simulated data.

**Extended Data Figure 2.**
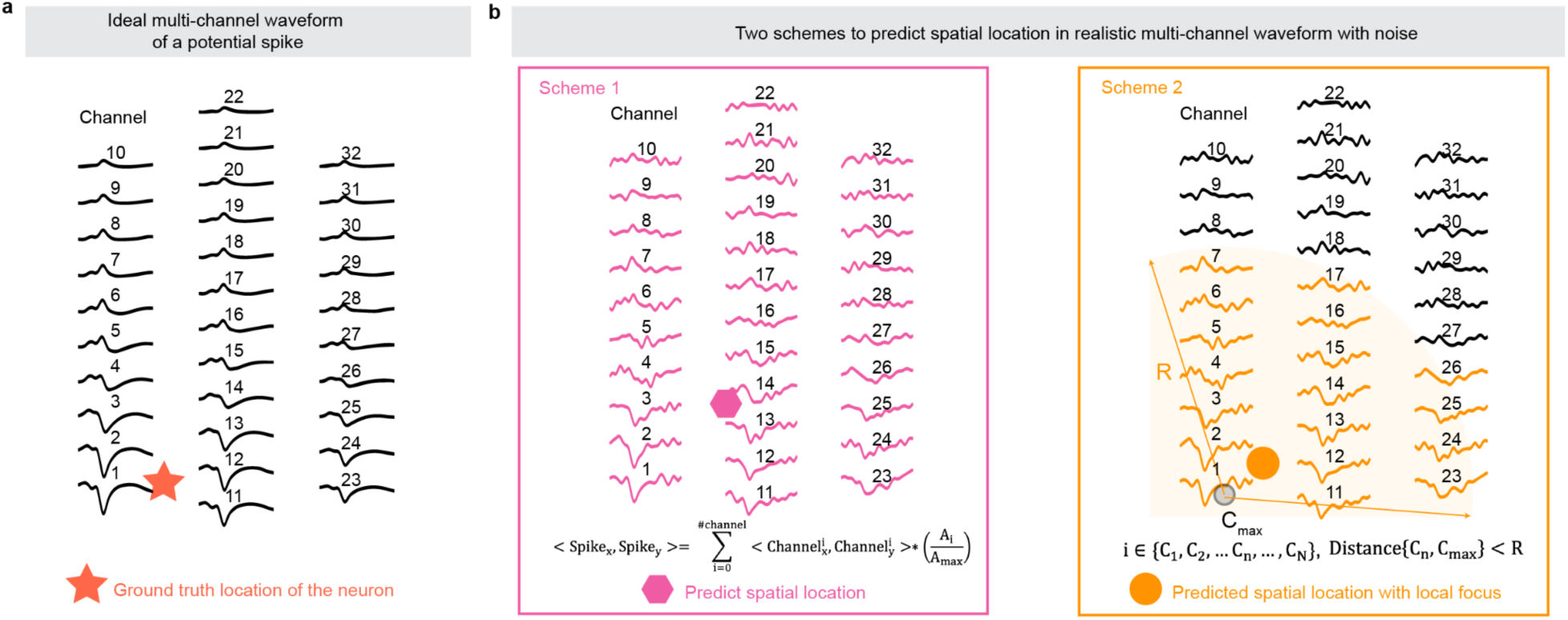
Comparison of spatial location prediction schemes for multi-channel electrophysiological data. **a,** Illustration that displays an ideal noise-free multi-channel waveform for a potential spike across 32 channels, highlighting the ground truth location of the neuron (indicated by the red star). **b,** Illustration of a predictive scheme for identifying the spatial location of spike activity in noisy multi-channel waveform data. The predicted location, highlighted in pink, utilizes a method based on the similarity between spike templates. **c,** Similar to (**b**), this panel presents a second scheme to predict the spatial location of spikes, highlighted in orange. This method focuses on local waveform characteristics to estimate the spike’s origin within a specified radius (R).

**Extended Data Figure 3.**
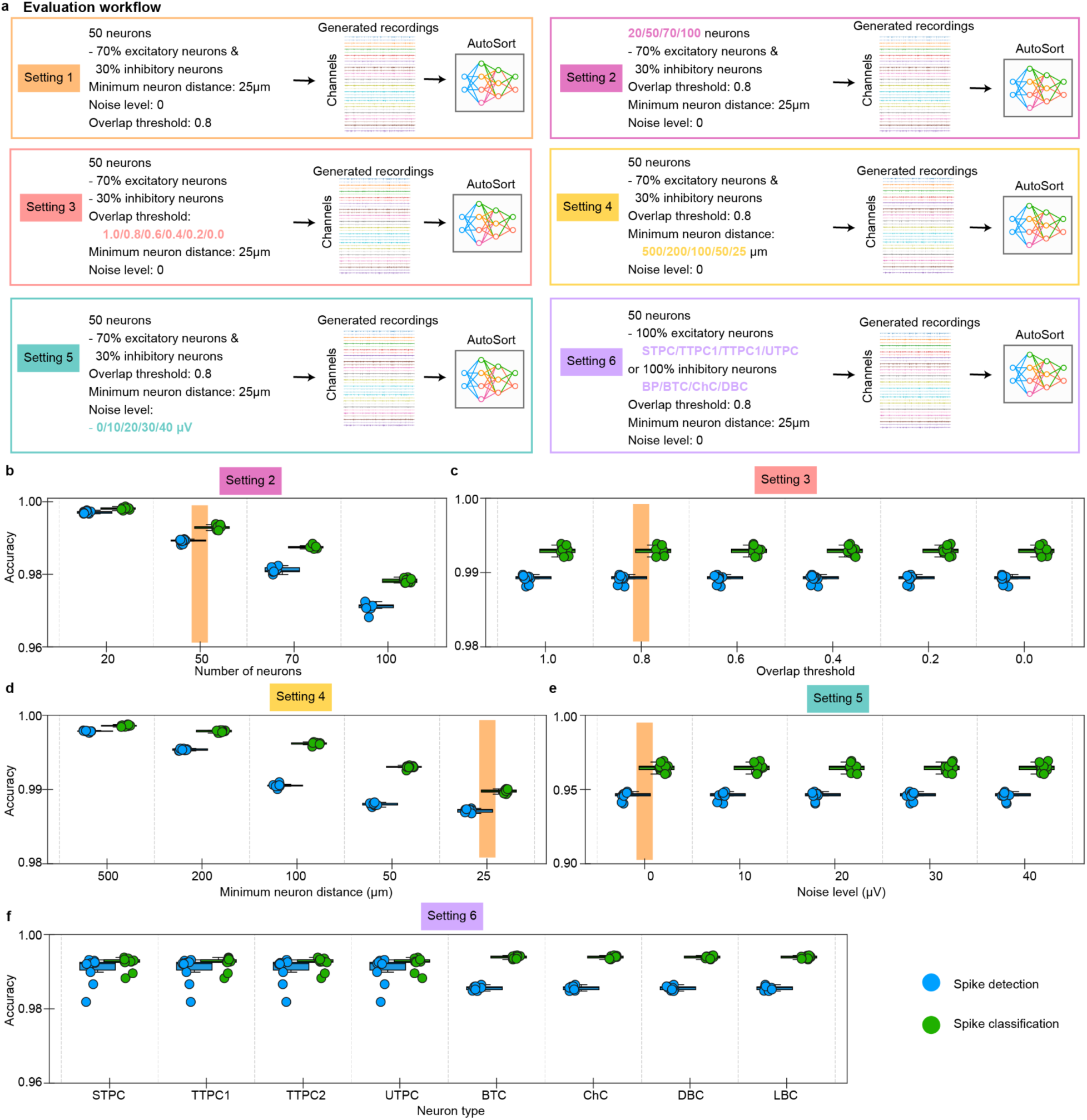
Additional combinations of experimentally relevant variables in simulated datasets for AutoSort performance evaluation. **a,** AutoSort evaluation workflow for different settings of the simulated dataset. The six settings include: Setting 1 (standard), corresponding to the conditions used in Figure 2, while Settings 2 through 6 modify one variable at a time while maintaining all others constant. Each setting is highlighted in distinct colors: Setting 2 adjusts the number of neurons from 20 to 100 (orange); Setting 3 alters the noise level from 0 to 40 µV (blue); Setting 4 expands the range of neuron counts from 20 to 1000 (pink); Setting 5 adjusts the minimum neuronal distance from 500 to 25 µm (yellow); and Setting 6 varies neuron types (purple). **b,** AutoSort’s accuracy across different settings, using ten models with varied random seeds to confirm result reliability. A vertical orange line indicates the parameters of Setting 1 from Figure 2. Dots represent the accuracy values for each AutoSort model. Box, 75% and 25% quantiles. Line, median. Whisker, the maxima/minima or to the median ± 1.5× inter quartile range (IQR).

**Extended Data Figure 4.**
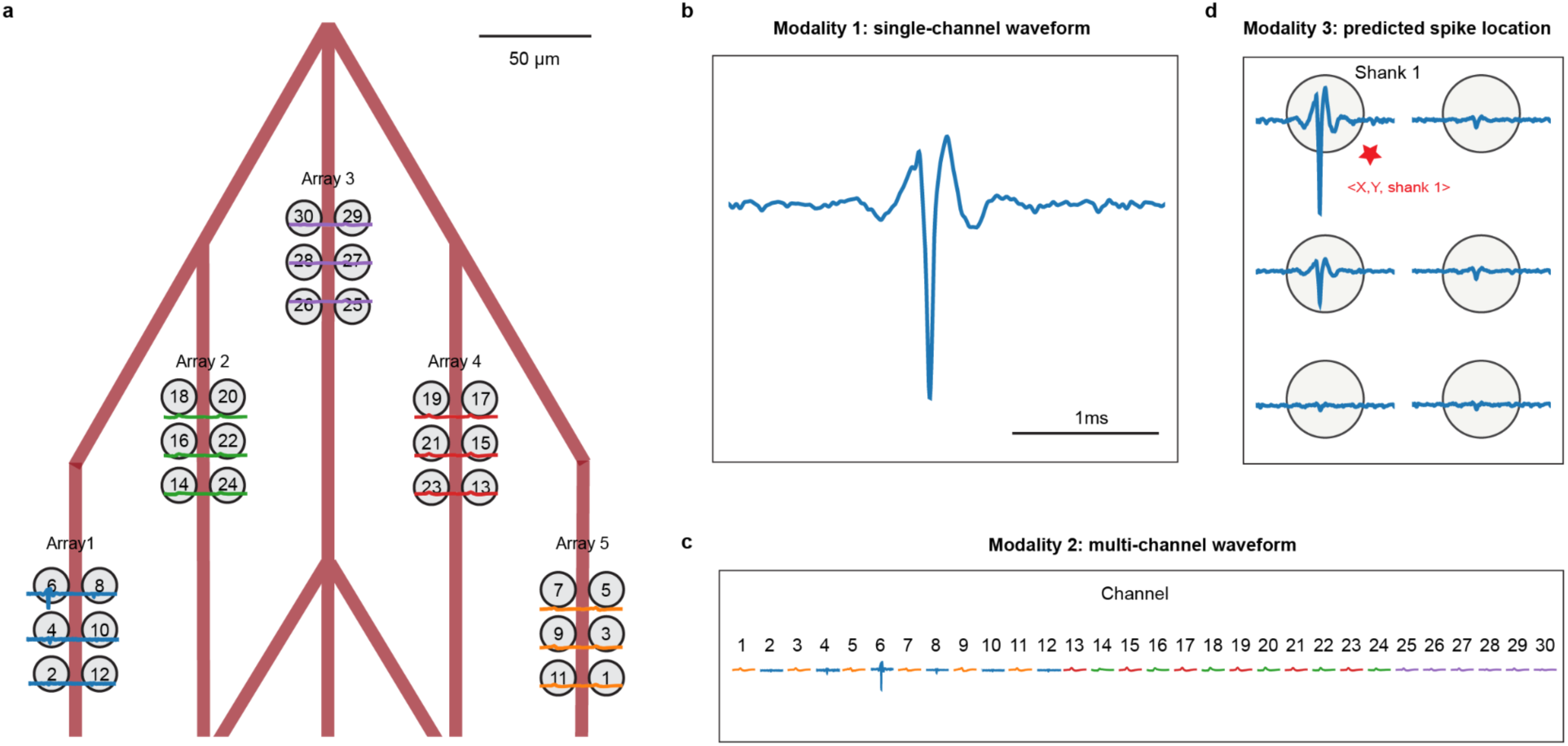
Representative tetrode-like electrode array multimodal on in vivo electrophysiological data. **a**, Illustration of the used mesh electronics with tetrode-like electrode arrays. Each mesh features five electrode arrays, with each array comprising six individually addressable electrodes for recording. This panel also includes representative waveforms from each of the channels. **b,** representative single-channel waveform from the channel with the highest amplitude within the array, showcasing the signal’s clarity and amplitude characteristics. **c,** Representative multi-channel waveform, displaying concatenated waveforms from all channels, providing a comprehensive view of the signal patterns across the entire array. **d,** Representative visualization of predicted spike locations. The display shows the first two components as the X and Y coordinates within a shank, while the third coordinate identifies the specific shank number where the spike is predicted to occur. The red star shows the inferred location calculated as detailed in Extended Data Figure 2.

**Extended Data Figure 5.**
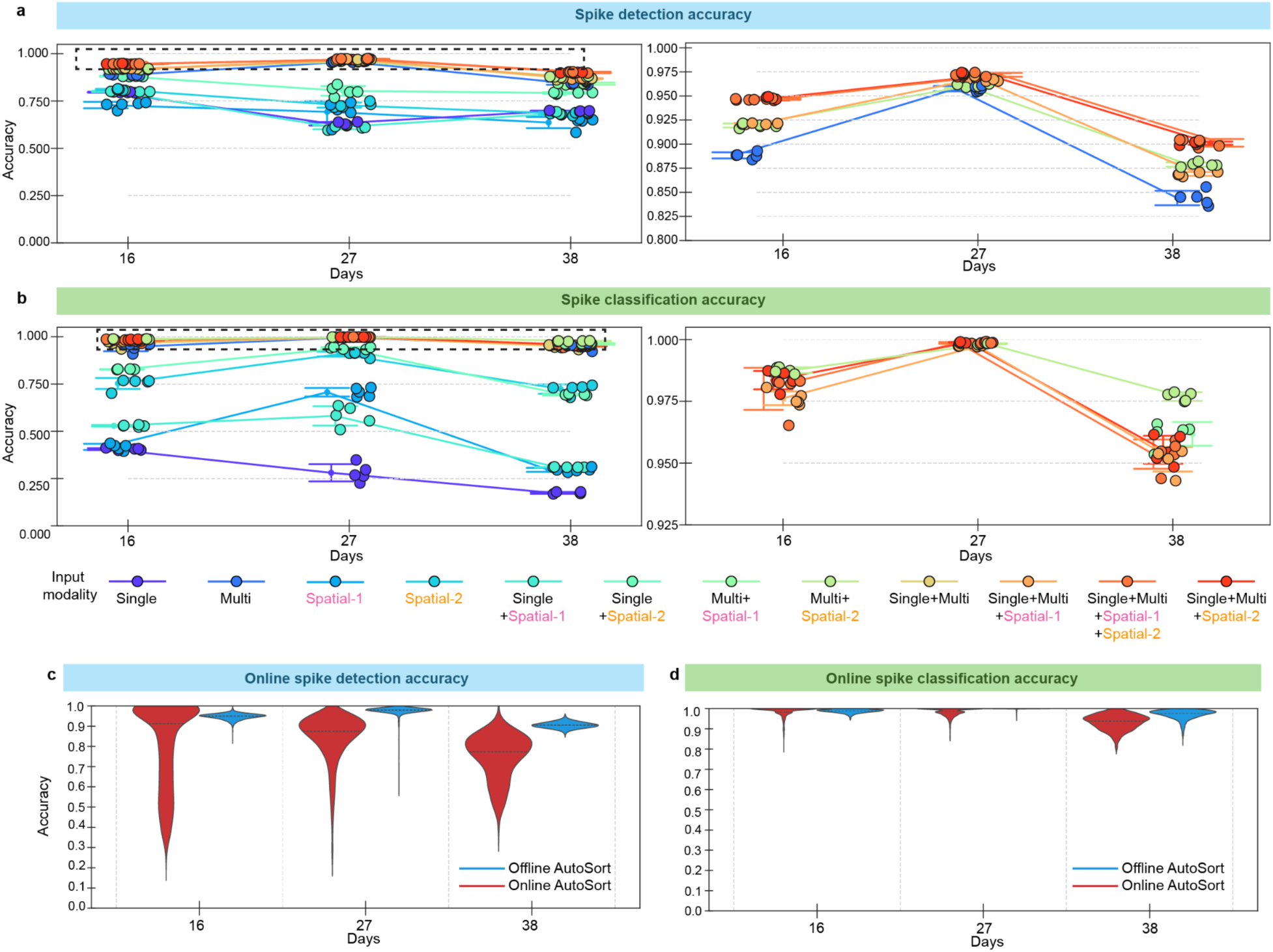
AutoSort accuracy on online long-term electrophysiological recording on mouse #2 dataset. **a,** Spike detection accuracy of AutoSort with different combinations of input modalities on representative chronic electrophysiological recording data, as in Figure 2. On the right, the highest-performing spike detection models are zoomed in for comparison. **b**, Same as in (a) but for the spike classification accuracy. **c,** Violin plots summarizing statistical spike detection accuracy between online (red) and offline (blue) AutoSort analysis on the representative chronic recording data in Mouse #2 from the same days as in (**a**) and (**b**). **d,** Same as in (**c**) with the spike classification. To train the different seeds (num = 5) for the models, we used day 11 (after 21 days post implantation) as the training dataset and tested on days 16, 27, and 38 (after 21 days post implantation) as representative data points. For all the strip plots, dots and line plots represent the mean accuracy values across the different seeds (num = 5) of the models for each test data point, and error bars indicate the SD. For all violin plots, the line inside the violin plots represents the median, providing an overview of the data accuracy distribution.

**Extended Data Figure 6.**
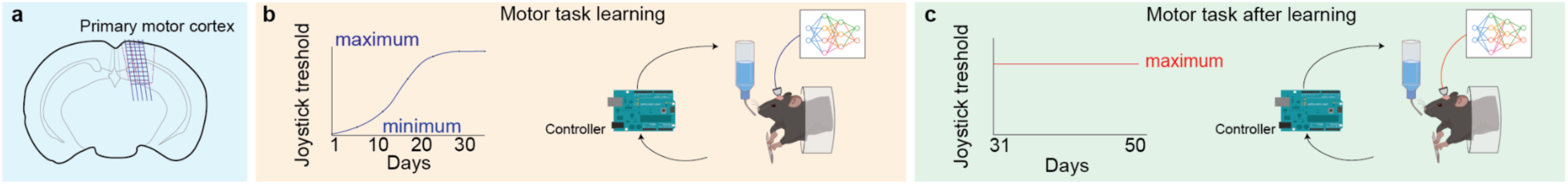
Overview of the animal experiments setup. **a,** Diagram showing the implantation regions within the motor cortex. **b,** Experimental timeline where animals learn a motor task by pulling a joystick to receive a water reward. The controller records joystick movement and delivers the reward when a certain threshold is reached. **c,** The level of proficiency is indicated by consistently achieving the joystick threshold.

## References

1. Lütcke, H., Margolis, D. J., & Helmchen, F. Steady or changing? Long-term monitoring of neuronal population activity. Trends Neurosci. 36, 375–384 (2013).

2. Stacey, W., & Litt, B. Technology Insight: neuroengineering and epilepsy—designing devices for seizure control. Nat Rev Neurol 4, 190–201 (2008).

3. Gilron, R. et al. Long-term wireless streaming of neural recordings for circuit discovery and adaptive stimulation in individuals with Parkinson’s disease. Nat. Biotechnol. 39, 1078–1085 (2021).

4. Tang, X., Shen, H., Zhao, S., Li, N., & Liu, J. Flexible brain-computer interfaces. Nat. Electron. 6, 109–118 (2023).

5. Tang, X., He, Y., & Liu, J. Soft bioelectronics for cardiac interfaces. Biophys. Rev. 3, 011301 (2022).

6. Luan, L. et al. Recent advances in electrical neural interface engineering: minimal invasiveness, longevity, and scalability. Neuron 108, 302–321 (2020).

7. Fu, T. M. et al. Stable long-term chronic brain mapping at the single-neuron level. Nat. Methods 13, 875–882 (2016).

8. Zhao, S. et al. Tracking neural activity from the same cells during the entire adult life of mice. Nat. Neurosci. 26, 696–710 (2023).

9. Zhao, Z. et al. Ultraflexible electrode arrays for months-long high-density electrophysiological mapping of thousands of neurons in rodents. *Nat*. Biomed. Eng. 7, 520–532 (2023).

10. Hong, G. & Lieber, C. M. Novel electrode technologies for neural recordings. Nat. Rev. Neurosci. 20, 330–345 (2019).

11. Zhou, W. et al. Long term stability of nanowire nanoelectronics in physiological environments. Nano Lett. 14, 1614–1619 (2014).

12. Fang, Y. et al. Dissecting biological and synthetic soft-hard interfaces for tissue-like systems. Chem. Rev. 122, 5233–5276 (2021).

13. Yasar, T. B. et al. Months-long tracking of neuronal ensembles spanning multiple brain areas with Ultra-Flexible Tentacle Electrodes. Nat. Commun. 15, 4822 (2024).

14. Guan, S. et al. Self-assembled ultraflexible probes for long-term neural recordings and neuromodulation. Nat. Protoc. 18, 1712–1744 (2023).

15. Liu, T. et al. A high-density 1,024-channel probe for brain-wide recordings in non-human primates. Nat. Neurosci. 27, 1620–1631 (2024).

16. Steinmetz, N. A. et al. Neuropixels 2.0: A miniaturized high-density probe for stable, long-term brain recordings. Science 372, eabf4588 (2021).

17. Luo, T. Z. et al. An approach for long-term, multi-probe Neuropixels recordings in unrestrained rats. eLife 9, e59716 (2020).

18. Eshraghian, J. K. et al. Training spiking neural networks using lessons from deep learning. Proc. IEEE 111, 1016–1054. (2023)

19. Chung, J. E. et al. A fully automated approach to spike sorting. Neuron 95, 1381–1394 (2017).

20. Pachitariu, M., Sridhar, S., Pennington, J., & Stringer, C. Spike sorting with Kilosort4. Nat. Methods 21, 914–921(2024).

21. Magland, J. F. & Barnett, A. H. Unimodal clustering using isotonic regression: ISO-SPLIT. Preprint at *arXiv* https://arxiv.org/abs/1508.04841 (2015).

22. Wang, P. K. et al. Low-latency single channel real-time neural spike sorting system based on template matching. PloS One 14, e0225138. (2019).

23. Vargas-Irwin, C., & Donoghue, J. P. Automated spike sorting using density grid contour clustering and subtractive waveform decomposition. J. Neurosci. Methods 164, 1–18. (2007).

24. Radmanesh, M., Rezaei, A. A., Jalili, M., Hashemi, A., & Goudarzi, M. M. Online spike sorting via deep contractive autoencoder. Neural Netw. 155, 39–49 (2022).

25. van Beest, E. H. et al. Tracking neurons across days with high-density probes. Nat Methods, 1–10 (2024)

26. Karkare, V., Gibson, S. & Markovic, D. A 130-µW, 64-channel neural spike-sorting DSP chip. IEEE J. Solid-State Circ. 46, 1214–1222 (2011).

27. Buccino, A. P., & Einevoll, G. T. Mearec: a fast and customizable testbench simulator for ground-truth extracellular spiking activity. Neuroinformatics 19, 185–204 (2021).

28. 28. McInnes, L. & Healy, J. UMAP: uniform manifold approximation and projection for dimension reduction. Preprint at arXiv https://arXiv.org/abs/1802.03426 (2018).

29. Buccino, A. P. et al. SpikeInterface, a unified framework for spike sorting. eLife 9, e61834 (2020).

30. Schoonover, C. E., Ohashi, S. N., Axel, R., & Fink, A. J. Representational drift in primary olfactory cortex. Nature 594, 541–546 (2021)

31. Zhao, S. et al. Realigning representational drift in mouse visual cortex by flexible brain-machine interfaces. Preprint at *BioRxiv* 10.1101/2024.05.23.595627 (2024).

32. Pehlevan, C., Hu, T., & Chklovskii, D. B. A hebbian/anti-hebbian neural network for linear subspace learning: A derivation from multidimensional scaling of streaming data. Neural Comput. 27, 1461–1495 (2015).

33. Todorova, S., Sadtler, P., Batista, A., Chase, S., & Ventura, V. To sort or not to sort: the impact of spike-sorting on neural decoding performance. J. Neural Eng. 11, 056005 (2014).

34. Davies, M. et al. Loihi: A neuromorphic manycore processor with on-chip learning. IEEE Micro 38, 82–99 (2018).

35. Modha, D. S. et al. Neural inference at the frontier of energy, space, and time. Science 382, 329–335 (2023).

36. Furber, S. B., Galluppi, F., Temple, S., & Plana, L. A. The spinnaker project. Proc. IEEE 102, 652–665 (2014).

37. Wang, H., Liu, W., Bocchieri, A., & Li, Y. Can multi-label classification networks know what they don’t know? Adv. Neural Inf. Proc. Sys. 34, 29074–29087 (2021).

38. Boussard, J., Varol, E., Lee, H. D., Dethe, N., & Paninski, L. Three-dimensional spike localization and improved motion correction for Neuropixels recordings. Adv. Neural Inf. Proc. Sys. 34, 22095–22105 (2021).

39. Garcia, S., Windolf, C., Boussard, J., Dichter, B., Buccino, A. P., & Yger, P. A modular implementation to handle and benchmark drift correction for high-density extracellular recordings. eNeuro 11 (2024).

40. Zhang, Yizi, et al. Towards a “universal translator” for neural dynamics at single-cell, single-spike resolution. Preprint at arXiv 10.48550/arXiv.2407.14668 (2024).

